# Nucleocapsid-specific humoral responses improve the control of SARS-CoV-2

**DOI:** 10.1101/2022.03.09.483635

**Authors:** Tanushree Dangi, Sarah Sanchez, Mincheol Park, Jacob Class, Michelle Richner, Justin M. Richner, Pablo Penaloza-MacMaster

## Abstract

The spike protein of SARS-CoV-2 is a critical antigen present in all approved SARS-CoV-2 vaccines. This surface viral protein is also the target for all monoclonal antibody therapies, but it is unclear whether antibodies targeting other viral proteins can also improve protection against COVID-19. Here, we interrogate whether nucleocapsid-specific antibodies can improve protection against SARS-CoV-2. We first immunized mice with a nucleocapsid-based vaccine, and then transferred sera from these mice into naïve mice. On the next day, the recipient mice were challenged intranasally with SARS-CoV-2 to evaluate whether nucleocapsid-specific humoral responses affect viral control. Interestingly, mice that received nucleocapsid-specific sera exhibited enhanced control of a SARS-CoV-2 infection. These findings provide the first demonstration that humoral responses specific to an internal coronavirus protein can help clear infection, warranting the inclusion of other viral antigens in next-generation SARS-CoV-2 vaccines and providing a rationale for the clinical evaluation of nucleocapsid-specific monoclonals to treat COVID-19.

**Highlights:** A SARS-CoV-2 nucleocapsid vaccine elicits robust nucleocapsid-specific antibody responses.

This nucleocapsid vaccine generates memory B cells (MBC).

Nucleocapsid-specific humoral responses do not prevent SARS-CoV-2 infection.

Nucleocapsid-specific humoral responses help control a SARS-CoV-2 infection.

## Introduction

Severe acute respiratory syndrome coronavirus 2 (SARS-CoV-2) has infected more than 400 million people and continues to spread around the globe. Although vaccines and monoclonal antibody therapies can prevent severe disease and death, breakthrough infections do occur, highlighting the need for improving current vaccines and antibody therapies (Baden et al., 2021; Keehner et al., 2021; Levine-Tiefenbrun et al., 2021; Logunov et al., 2021; Polack et al., 2020; Sadoff et al., 2021; Tande et al., 2021; Voysey et al., 2021; Zhu et al., 2020). The SARS-CoV-2 spike protein is critical for viral entry, and thus, this protein is the main antigen present in current SARS-CoV-2 vaccines. However, it is unclear if antibody responses elicited by other viral antigens can contribute to host protection. In particular, it is unknown if antibodies specific to internal viral proteins, such as the nucleocapsid protein which does not play a role in viral entry, could confer protection against SARS-CoV-2. Knowing if non-spike-specific antibodies are protective could facilitate the development of more potent vaccines and monoclonal antibody cocktails for coronavirus infections. In this study, we interrogated whether humoral responses elicited by a nucleocapsid-based vaccine could confer protection against a SARS-CoV-2 challenge in K18-hACE2 mice. Interestingly, we show that nucleocapsid-specific humoral responses help control SARS-CoV-2 infection.

## Results

### Immunogenicity of a nucleocapsid-based vaccine regimen

We first primed C57BL/6 mice intramuscularly with an adenovirus serotype 5 vector expressing SARS-CoV-2 nucleocapsid (Ad5-N) (Dangi et al., 2021a; Dangi et al., 2021b; Joag et al., 2021) at a dose of 10^11^ PFU per mouse, followed by booster with 100 μg of nucleocapsid protein three weeks later. As controls, we immunized mice with an “empty” Ad5 vector (Ad5-E) followed by a PBS boost. After 2 weeks post-boost, we measured nucleocapsid-specific immune responses (**Figure 1A**). Mice immunized with the nucleocapsid vaccine regimen exhibited robust nucleocapsid-specific CD8 T cell responses (**Figure 1B**), memory B cell responses (**Figure 1C**), and antibody responses (**Figure 1D**).

**Figure 1.**
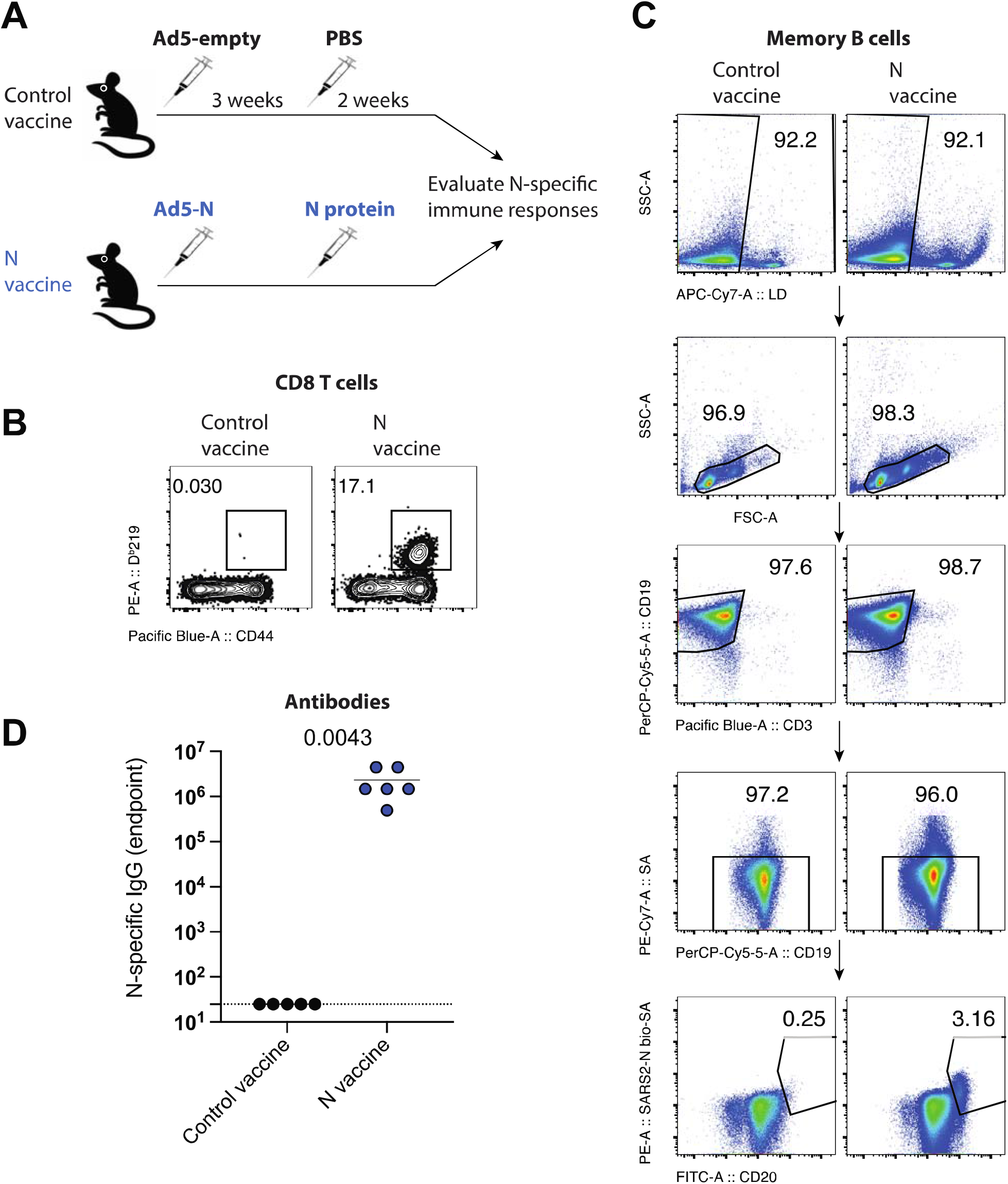
Immunogenicity of a SARS-CoV-2 nucleocapsid vaccine regimen. **(A)** Experimental approach for evaluating immune responses after nucleocapsid vaccination. **(B)** Representative FACS plots showing the frequencies of SARS-CoV-2 nucleocapsid-specific CD8 T cells (D^b^N219+) in PBMCs. **(C)** Representative FACS plots showing the frequencies of SARS-CoV-2 nucleocapsid-specific memory B cells in spleen. Splenocytes were MACS-purified by negative selection to enrich for B cells, facilitating visualization of nucleocapsid-specific B cells. **(D)** Summary of SARS-CoV-2 nucleocapsid-specific antibodies in sera. Data are from week 2 post-boost, and from an experiment with n=5-6 per group. Experiment was performed twice with similar results. Indicated P-values were calculated using Mann Whitney test.

Prior studies have suggested that nucleocapsid-specific T cells can help clear SARS-CoV-2 infection (Dangi *et al*., 2021a; Dangi *et al*., 2021b; Joag *et al*., 2021; Matchett et al., 2021). However, it is still unclear if nucleocapsid-specific antibodies play any role in antiviral control, since the nucleocapsid is an internal viral protein that is not involved in viral entry. We performed focus reduction neutralization titer (FRNT) assays using live SARS-CoV-2 to examine if nucleocapsid-specific antibodies prevent SARS-CoV-2 infection (**Figure 2A**). We used live SARS-CoV-2 instead of a pseudovirus, because the former contains all viral proteins, including nucleocapsid. As positive control, we used sera from mice that received a spike-based adenovirus vaccine (Ad5-S, similar to the CanSino vaccine and the Sputnik vaccine). As expected (Dangi *et al*., 2021b; Sanchez et al., 2021), sera from mice that received this spike-based vaccine prevented SARS-CoV-2 infection, even when the sera were diluted 450-fold (**Figure 2B**). However, sera from mice that received the nucleocapsid-based vaccine did not exert any antiviral effect in this in vitro infection assay (**Figure 2B–2C**). Taken together, only spike-specific antibodies can block SARS-CoV-2 infection, consistent with the widely established notion that these antibodies are protective, as they can block the first step in the SARS-CoV-2 life cycle (entry into host cells).

**Figure 2.**
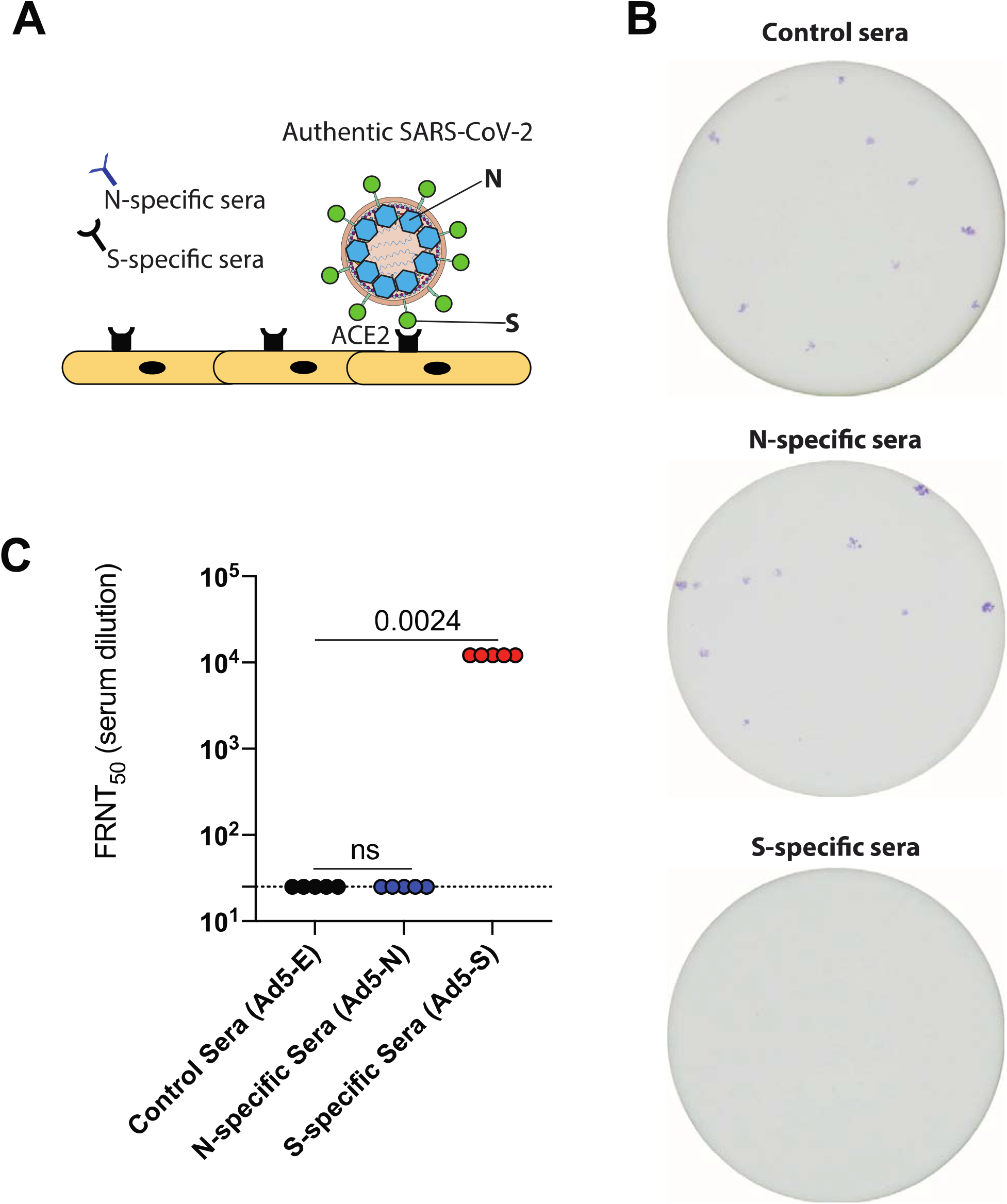
Nucleocapsid-specific humoral responses do not prevent SARS-CoV-2 infection. **(A)** Experimental approach for performing focus reduction neutralization titer (FRNT) assays on Vero cells using live SARS-CoV-2 (USA-WA1/2020). See Materials and Methods for technical information. **(B)** Representative wells showing the frequencies of SARS-CoV-2+ cells (1:450 sera dilution). **(C)** Summary of FRNT_50_ titers in sera. Data are from week 3 post-vaccination. Data are from an experiment with n=5 per group. Experiment was performed twice with similar results. Indicated P-values were calculated using Kruskal-Wallis test (multiple comparisons).

### Nucleocapsid-specific humoral responses help clear a SARS-CoV-2 infection in vivo

Antibody responses exert antiviral functions by various mechanisms, including viral neutralization and effector mechanisms. Although nucleocapsid-specific sera did not prevent SARS-CoV-2 infection in vitro, we reasoned that it could confer protection in vivo via other mechanisms different than neutralization. To test this hypothesis, we performed a passive immunization study to evaluate whether the specific transfer of nucleocapsid-specific antibodies can confer a clinical benefit (**Figure 3A**). We adoptively transferred 500 μL of nucleocapsid-specific sera into naïve K18-hACE2 mice, which are susceptible to SARS-CoV-2 (Bao et al., 2020; Dangi *et al*., 2021a; Oladunni et al., 2020; Rosenfeld et al., 2021; Winkler et al., 2020; Zheng et al., 2021). One day after sera transfer, we challenged these K18-hACE2 mice intranasally with 10^3^ PFU of SARS-CoV-2 (isolate USA-WA1/2020), and at day 4 post-challenge we quantified viral loads in lungs by RT-qPCR. Interestingly, the mice that received nucleocapsid-immune sera showed a significant reduction in viral titers relative to control (**Figure 3B**). These findings suggest that even though nucleocapsid-specific antibodies do not exert any antiviral effect in vitro, they can exert antiviral effects in vivo.

**Figure 3.**
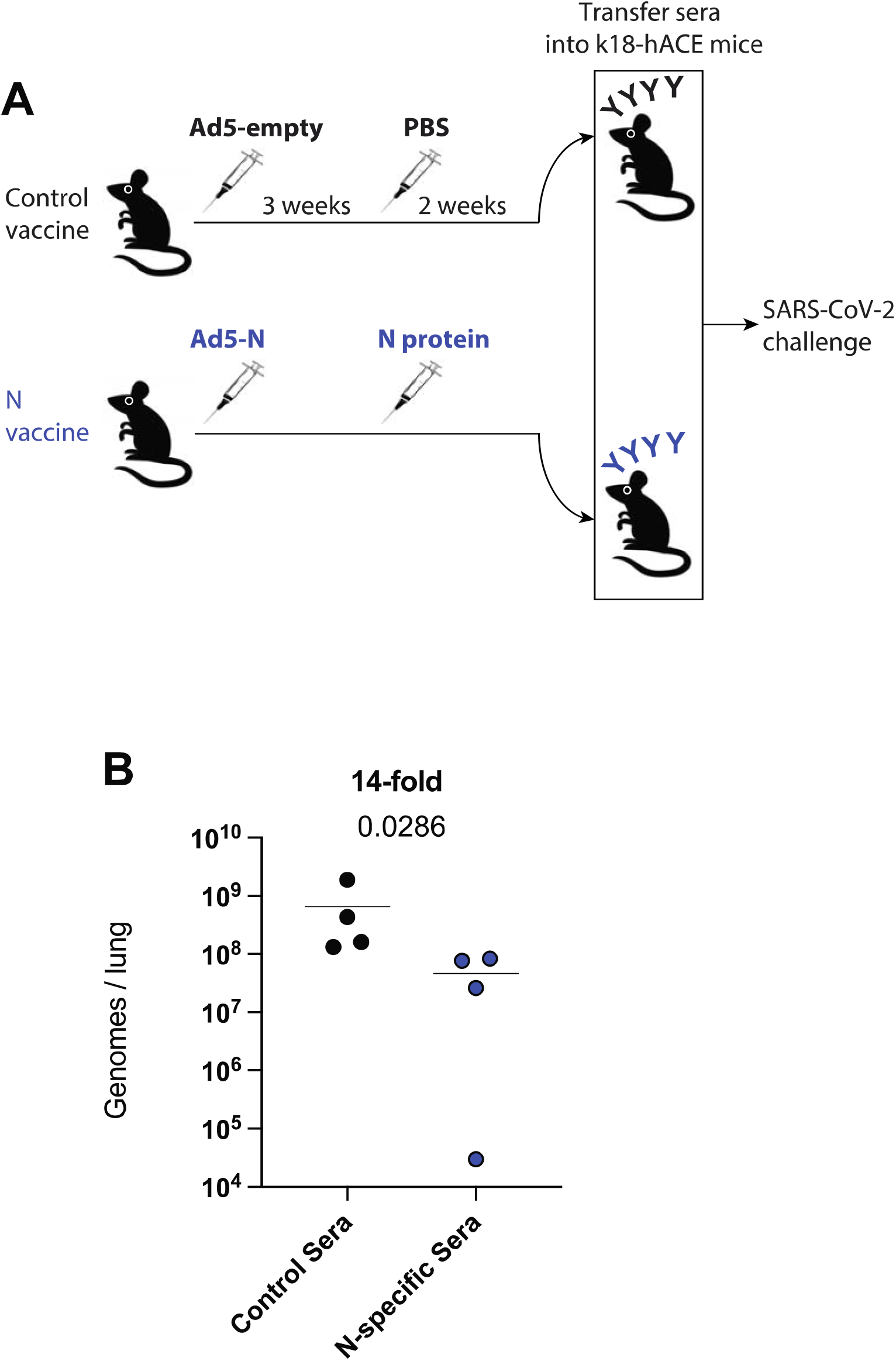
Nucleocapsid-specific humoral responses improve the control of SARS-CoV-2 infection. C57BL/6 mice were primed intramuscularly with Ad5 expressing nucleocapsid (Ad5-N), and after 3 weeks they were boosted with soluble nucleocapsid (N) protein. After 2 weeks post-boost, sera from these mice were pooled and 500 μL of these sera were transferred into naïve K18-hACE2 recipient mice. On the following day, the K18-hACE2 mice were challenged intranasally with 10^3^ PFU of SARS-CoV-2. RNA was harvested from the lungs at day 4 post-infection, and viral RNA was quantified by RT-qPCR. Challenges were performed with a total of 4 mice per group in Biosafety level 3 (BSL-3) facilities. Indicated P-values were calculated using Mann Whitney test.

## Discussion

The SARS-CoV-2 spike protein mediates viral entry by binding to the ACE2 receptor, and therefore, this protein is the most important target for vaccines and monoclonal antibody therapies. All approved SARS-CoV-2 vaccines target the spike protein with the goal of generating spike-specific antibodies that prevent viral entry. However, it is unclear if antibodies of other specificities (e.g. internal viral proteins that do not mediate viral entry) can play a role in antiviral protection. In particular, nucleocapsid-specific antibodies are generated after SARS-CoV-2 infection or after immunization with experimental nucleocapsid-based vaccines (Dangi *et al*., 2021a; Dangi *et al*., 2021b), but it is unclear if nucleocapsid-specific antibody responses can protect the host against a SARS-CoV-2 infection.

To the best of our knowledge, the data presented here provide the first demonstration that nucleocapsid-specific humoral responses can help clear a SARS-CoV-2 infection. These findings may be important for the development of next-generation SARS-CoV-2 vaccines and for developing novel antibody therapies. Current monoclonal antibody therapies for COVID-19 target only the spike protein, and many of these therapies have lost efficacy against variants, since the spike protein is highly variable. Our data provide a rationale for the clinical evaluation of monoclonal antibody therapies targeting the nucleocapsid protein, which is relatively conserved among coronaviruses. Although nucleocapsid-specific antibodies do not prevent infection with SARS-CoV-2, they could help control infection. It is possible that nucleocapsid-specific antibodies may also synergize with spike-specific antibodies in current monoclonal therapies.

In a prior paper we demonstrated that a nucleocapsid-based vaccine confers limited protection against a SARS-CoV-2 challenge when given as a “single vaccine” without a spike-based vaccine. At first glance, this prior study appears to contradict our results, but in the prior study we used a high challenge dose of SARS-CoV-2 (5×10^4^ PFU), which could have overwhelmed nucleocapsid-specific immunity. In the present study, however, we used a more physiological challenge dose of SARS-CoV-2 (10^3^ PFU), and we also evaluated viral loads at a later time post-infection. Thus, it is thus possible that the antiviral effects of nucleocapsid-specific antibodies vary depending on the virus inoculum and the time of viral infection.

For more than a century, convalescent plasma therapy has been used to treat infectious diseases. Convalescent plasma therapy has shown efficacy against COVID-19, but its benefits are more apparent in patients who receive plasma with high antibody titers, which can only be obtained from convalescent donors early after infection (Casadevall et al., 2021; Jabal et al., 2021). A limitation of our study is that we only performed adoptive sera transfers early after nucleocapsid vaccination. It is unknown if a similar level of protection would be observed if the passive immunizations were performed using sera from later time points post-vaccination. We detected memory B cells, suggesting that the nucleocapsid-specific antibody response was long-lived, but future studies are needed to evaluate the durability of nucleocapsid-specific antibody responses and the mechanisms by which they clear SARS-CoV-2 infection. Overall, we show that passive immunization using nucleocapsid-specific sera improves the control of a SARS-CoV-2 infection. These data provide insights for next-generation vaccines and for expanding the armamentarium of antibody-based therapies for COVID-19.

## STAR Methods

### Key resources table

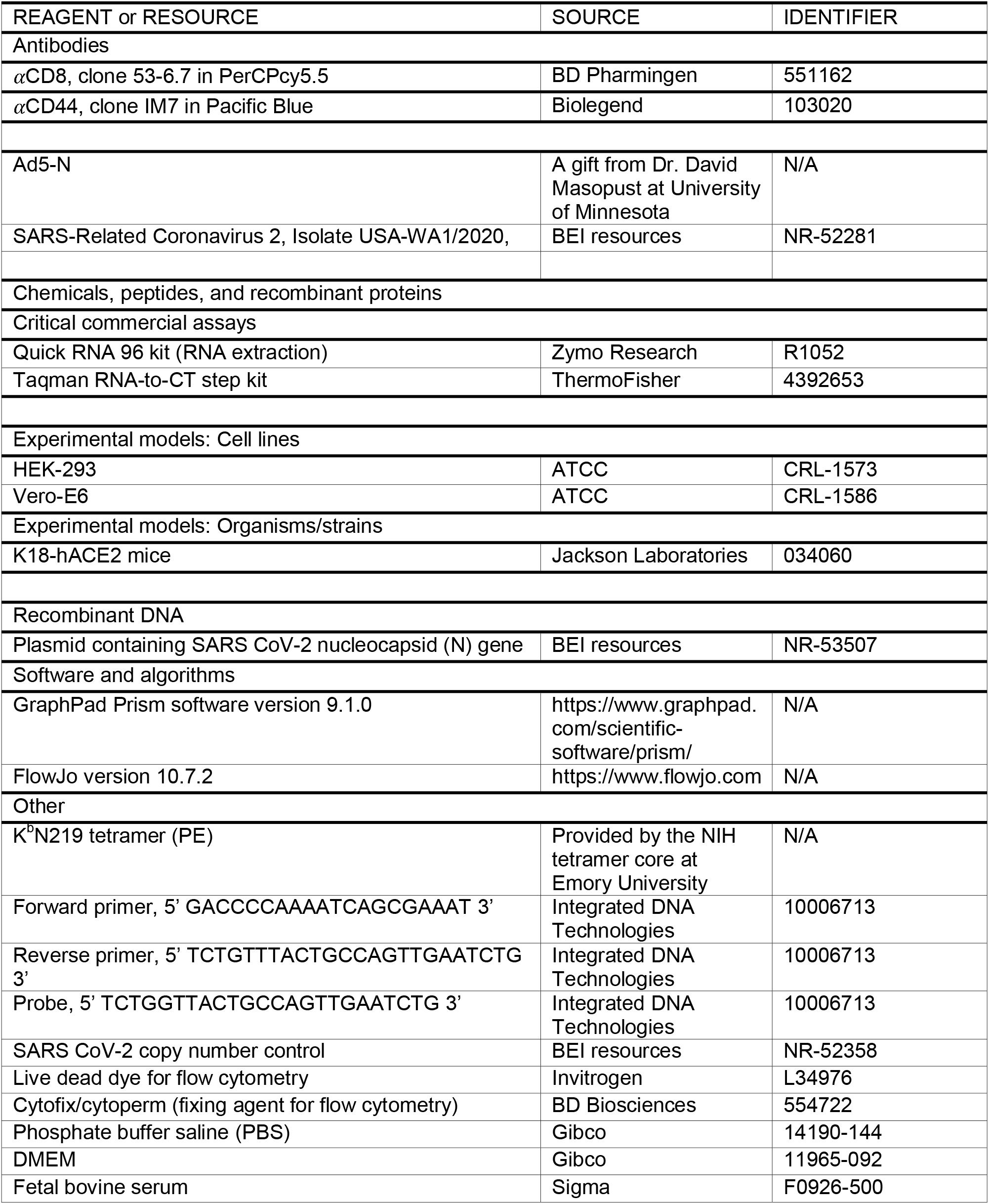

### Resource availability

#### Lead contacts

Further information and requests for resources and reagents should be directed to and will be fulfilled by the lead contact Pablo Penaloza-MacMaster and/or Justin Richner

### Materials availability

For Ad5-SARS CoV-2 nucleocapsid viral vector access, contact David Masopust at.

### Data and code availability

This article does not report original code. The article includes all the data sets and analyses generated for this study. Any additional information required to reanalyze the data reported in this paper is available from the lead contact upon request.

### Experimental models and subject details

#### Animals and ethics statement

Mice were purchased from Jackson laboratories and were housed at Northwestern University or University of Illinois at Chicago (UIC) animal facility. All procedures were performed with the approval of the center for comparative medicine at Northwestern University and the UIC IACUC. Adult mice, approximately half females and half males were used for the immunogenicity experiments included in this study. For the challenge studies, female mice were used.

### Method Details

#### Cell lines

Adenoviral vectors were propagated using HEK293 cells purchased from ATCC (cat # CRL-1573). Vero E6 cells were used to propagate SARS CoV-2 isolate USA-WA1/2020 (BEI resources, NR-52281). Cells were not authenticated as they were purchased from a reputable vendor and certificate of analysis was obtained.

#### Mice and vaccinations

6-8-week-old K18-hACE2 mice were used. These mice express the human ACE2 protein behind the keratin 18 promoter, directing expression in epithelial cells. Mice were purchased from Jackson laboratories (Stock No: 034860). Mice were immunized intramuscularly (50 μL per quadriceps) with an Ad5 vector expressing SARS-CoV-2 nucleocapsid protein (Ad5-N) at 10^11^ PFU per mouse, and N protein; diluted in sterile PBS. Ad5-N was a kind gift of the Masopust/Vezys laboratory (Joag *et al*., 2021). This is a non-replicating Ad5 vector (E1/E3 deleted). The vector contains a CMV (Cytomegalovirus) promoter driving the expression of the respective proteins. The Ad5 vector was propagated on trans-complementing HEK293 cells (ATCC), purified by cesium chloride density gradient centrifugation, titrated, and then frozen at −80□°C.

#### SARS-CoV-2 virus and infections

SARS-Related Coronavirus 2, Isolate USA-WA1/2020, NR-52281 was deposited by the Centers for Disease Control and Prevention and obtained through BEI Resources, NIAID, NIH. Virus was propagated and tittered on Vero-E6 cells (ATCC). In brief, Vero cells were passaged in DMEM with 10% Fetal bovine serum (FBS) and Glutamax. Cells less than 20 passages were used for all studies. Virus stocks were expanded in Vero-E6 cells following a low MOI (0.01) inoculation and harvested after 4 days. Viral titers were determined by plaque assay on Vero-E6 cells. Viral stocks were used after a single expansion (passage = 1) to prevent genetic drift.

K18-hACE2 mice were anesthetized with isoflurane and challenged with 10^3^ PFU of SARS-CoV-2 intranasally. Mouse infectious were performed at the University of Illinois at Chicago (UIC) following BL3 guidelines with approval by the UIC Institutional Animal Care and Use Committee (IACUC).

#### SARS-CoV-2 quantification in lungs

Lungs were harvested from infected mice and homogenized in PBS. RNA was isolated with the Zymo 96-well RNA isolation kit (Catalog #: R1052) following the manufacturer’s protocol. SARS-CoV-2 viral burden was measured by RT-qPCR using Taqman primer and probe sets from IDT with the following sequences: Forward 5’ GAC CCC AAA ATC AGC GAA AT 3’, Reverse 5’ TCT GGT TAC TGC CAG TTG AAT CTG 3’, Probe 5’ ACC CCG CAT TAC GTT TGG TGG ACC 3’. A SARS-CoV-2 copy number control was obtained from BEI (NR-52358) and used to quantify SARS-CoV-2 genomes.

#### Focus Reduction Neutralization Titer (FRNT) Assay using live SARS-CoV-2

FRNT assays were performed by doing serial dilutions of heat-inactivated serum from vaccinated mice, incubated with live SARS-CoV-2 (isolate USA-WA1/2020) for one hour at 37°C before infecting a monolayer of Vero cells in a 96 well plate. One hour after infection, cells were overlaid with 1% (w/v) methylcellulose in 2% FBS, 1X MEM. Plates were fixed for 30 minutes with 4% PFA 24 hours after infection. Staining involved 1° anti-SARS guinea pig (1:15,000, NR-10361 from BEI Resources) and 2° goat anti-guinea pig HRP (200 ng/ml) in Perm Wash Buffer (0.1% Saponin, 0.1% BSA, in PBS). Treatment with TrueBlue peroxidase substrate (KPL) produced FFU that were quantified with an ImmunoSpot^®^ ELISpot plate scanner (Cellular Technology Limited).

#### Reagents, flow cytometry and equipment

Single cell suspensions were obtained from PBMCs or tissues. Dead cells were gated out using Live/Dead fixable dead cell stain (Invitrogen). SARS-CoV-2 nucleocapsid protein was biotinylated and conjugated to streptavidin-PE for detection of nucleocapsid-specific memory B cells on MACS-purified B cells. MHC class I monomers (K^b^219, LALLLLDRL) were used for detecting virus-specific CD8 T cells, and were obtained from the NIH tetramer facility located at Emory University. MHC monomers were tetramerized in-house. Cells were stained with fluorescently-labeled antibodies against CD8α (53-6.7 on PerCP-Cy5.5), CD44 (IM7 on Pacific Blue), and K^b^N219 (PE). Fluorescently-labeled antibodies were purchased from BD Pharmingen, except for anti-CD44 (which was from Biolegend). Flow cytometry samples were acquired with a Becton Dickinson Canto II or an LSRII and analyzed using FlowJo v10 (Treestar).

#### SARS-CoV-2 nucleocapsid specific ELISA

Binding antibody titers were quantified using ELISA as described previously (Dangi et al., 2020; Palacio et al., 2020), using nucleocapsid protein as coating antigens. In brief, 96-well flat bottom plates MaxiSorp (Thermo Scientific) were coated with SARS-CoV-2 nucleocapsid protein, washed and blocked. Serial sera dilutions were performed. Absorbance was measured using a Spectramax Plus 384 (Molecular Devices).

#### Quantification and statistical analysis

Statistical analyses are indicated on the figure legend. Dashed lines in data figures represent limit of detection. Statistical significance was established at p ≤0.05 and was generally assessed by Mann Whitney tests, unless indicated otherwise in figure legends. Data were analyzed using Prism (Graphpad).

## Author Contributions

T.D., M.P., and S.S. performed the vaccination and immunogenicity experiments, and expressed the nucleocapsid protein that was used for the vaccine and ELISA assays. J.C. performed the challenge experiments. P.P.M. and J.R. designed the experiments and secured funding. P.P.M. wrote the paper, with feedback from all authors.

## Acknowledgements

We thank Dr. Thomas Gallagher, David Masopust and Vaiva Vezys for discussions and reagents. This work was possible with a grant (R01AI150672) to J.M.R.; a grant from the National Institute of Biomedical Imaging and Bioengineering (NIBIB U54 EB027049) to P.P.M.; and a grant from the National Institute on Drug Abuse (NIDA, DP2DA051912) to P.P.M.

## Declaration of Interests

The authors declare that no conflicts of interests exist.

## Notes

### Competing Interest Statement

The authors have declared no competing interest.

